# A novel rare c. -39C>T mutation in the *PROS1* 5’UTR causing PS deficiency by creating a new upstream translation initiation codon and inhibiting the production of the natural protein

**DOI:** 10.1101/2020.03.28.007328

**Authors:** Sylvie Labrouche-Colomer, Omar Soukarieh, Carole Proust, Christine Mouton, Yoann Huguenin, Maguelonne Roux, Céline Besse, Anne Boland, Robert Olaso, Joël Constans, Jean-François Deleuze, Pierre-Emmanuel Morange, Béatrice Jaspard-Vinassa, David-Alexandre Trégouët, on behalf of the GenMed consortium

**Affiliations:** CHU de Bordeaux, Laboratoire d’Hématologie, Pessac, France; INSERM UMR 1034, Biology of Cardiovascular Disease, University of Bordeaux, Pessac, France; INSERM UMR 1219, Bordeaux Population Health Research Center, University of Bordeaux, Bordeaux, France; CHU de Bordeaux, Service de pédiatrie médicale, Bordeaux, France; Human Evolutionary Genetics Unit, Institut Pasteur, UMR2000, CNRS, Paris 75015, France; Université Paris-Saclay, CEA, Centre National de Recherche en Génomique Humaine, 91057, Evry, France; Centre d’Etude du Polymorphisme Humain, Fondation Jean Dausset, Paris, France; C2VN INSERM UMR 1263, INRA, Aix-Marseille University, Marseille, France

**Author notes:** These two authors equally contributed to the work. ^#^Correspondence to: Dr David-Alexandre Trégouët, INSERM U1219, Bordeaux Population Health Research Center, 33076 Bordeaux, France.

**Keywords:** Protein S deficiency, venous thrombosis, whole genome sequencing, Open Reading Frame, mutation

## Abstract

Inherited Protein S deficiency (PSD) (MIM176880) is a rare automosal dominant disorder caused by rare mutations, mainly located in the coding sequence of the structural *PROS1* gene, and associated with an increased risk of venous thromboembolism. To identify the molecular defect underlying PSD observed in an extended French pedigree with 7 PSD affected members in who no candidate deleterious *PROS1* mutation was detected by Sanger sequencing of *PROS1* exons and their flanking intronic regions or via a MLPA approach, a whole genome sequencing strategy was adopted. This led to the identification of a never reported C to T substitution at c.-39 from the natural ATG codon of the *PROS1* gene that completely segregates with PSD in the whole family. This substitution ACG->ATG creates a new start codon upstream of the main ATG. We experimentally demonstrated that the variant generates a novel overlapping ORF and inhibits the translation of the wild type protein from the main ORF in HeLa cells. This work describes the first example of 5’UTR *PROS1* mutation causing PSD through the creation of an upstream ORF, a mutation that is not predicted to be deleterious by standard annotation softwares.

## INTRODUCTION

Protein S (PS) is, with Protein C (PC) and Antithrombin (AT), one the three main natural inhibitors of the coagulation cascade and plays thus a key role in the control of blood clot formation. PS is a vitamin-K dependent glycoprotein that acts as a cofactor for PC to inactivate factors Va (FVa) and VIIIa (FVIIIa) and to limit thrombin generation via direct interactions with factor Xa (FXa) and FVa (Castoldi & Hackeng, 2008). In human plasma, PS circulates both under a free and active form (∼40%) and an inactive form (∼60%) when complexed with C4b-binding protein. Generally, PS plasma concentration can be characterized by antigen measurements of the free and total PS levels or by PS activity. This led to the definition of three clinical subtypes of PS deficiencies: i) Type I refers to deficiency of both free and total PS as well as decreased PS activity, ii) Type II is defined by normal plasma levels but decreased PS activity while iii) Type III shows decreased free PS plasma levels and decreased PS activity but normal total PS plasma levels. Type I and Type III PS deficiencies account for ∼95% of PS deficiencies and are considered to be the heterogeneous clinical expression of the same molecular defect (Zöller *et al*, 1995).

Inherited PS deficiency (MIM176880) (PSD) is an autosomal dominant disorder caused by private or rare mutations in the structural *PROS1* gene. Previous works have shown that complete or partial PSD was associated with increased risk of venous thrombosis (VT) (Alhenc-Gelas *et al*, 2016, 2010; Lijfering *et al*, 2009; Pegelow *et al*, 1992; Pintao *et al*, 2013). The majority of PSD causing *PROS1* mutations are located in the coding regions of the gene or in their flanking sequences and are mainly missense or nonsense single point variations (Stenson *et al*, 2017). Some intronic splice variants (Choi *et al*, 2011; Menezes *et al*, 2017; Wang *et al*, 2019) and structural variants (Hurtado *et al*, 2009; Lind-Halldén *et al*, 2012; Seo *et al*, 2014) have also been reported. Note that, compared to exonic variations, very few variations have been described in the promoter region of the *PROS1* gene (Espinosa-Parrilla *et al*, 2000; Hurtado *et al*, 2008; Li *et al*, 2019; Sanda *et al*, 2007; Tang *et al*, 2013) and even fewer have been functionally characterized. To our knowledge, the c.-168C>T is the sole *PROS1* promoter variation that has been experimentally demonstrated to cause inherited PSD by affecting the core binding site of Sp1 transcription factor (Sanda *et al*, 2007).

We here describe a novel mutation in the 5’UTR region of the *PROS1* gene that causes PSD in a family with multiple relatives affected with VT. The mutation is a C to T substitution at c.-39 from the natural ATG codon and creates a novel ATG sequence upstream of the main open reading frame that alters the translation machinery of the protein.

## METHODS AND MATERIALS

### Recruitment of the family

The studied family was ascertained through a proband addressed to the specialized clinical hematology laboratory of the Pellegrin Hospital (Bordeaux) for a thrombophilia screening in the context of an objectively documented first VT episode early in life (20 years old). Standard thrombophilia screening revealed a Type I PSD (Fig 1; Supp Table 1). However, no candidate deleterious *PROS1* mutation nor large genomic rearrangement were detected by Sanger sequencing of the 15 *PROS1* exons and of their flanking intronic regions (Labrouche *et al*, 2003) or by multiplex ligation dependent probe amplification (MLPA) using Salsa MLPA kit (P112; MRC-Holland, Amsterdam, the Netherlands), respectively. As the proband reported at least one first degree relative with VT, family members were invited to participate to a genetic exploration upon informed consent agreement according to the Helsinki declaration. Inherited Type I or Type III PSD was diagnosed in 6 additional relatives (Fig 1), a seventh relative (individual IV-4) had very low PS activity but, without information on free and total PS levels, Type I, Type II or Type III PSD could not be objectively distinguished. All PSD patients with DNA available were negative for the presence of *PROS1* deleterious coding mutation and for the F5 Leiden (FV Arg506Gln, rs6025), the F2 G20210 (rs1799963) mutations and any other major biological risk factors for venous thrombosis including antithrombin-, PC-deficiency or anti-phospholipid antibodies.

**Figure 1:**
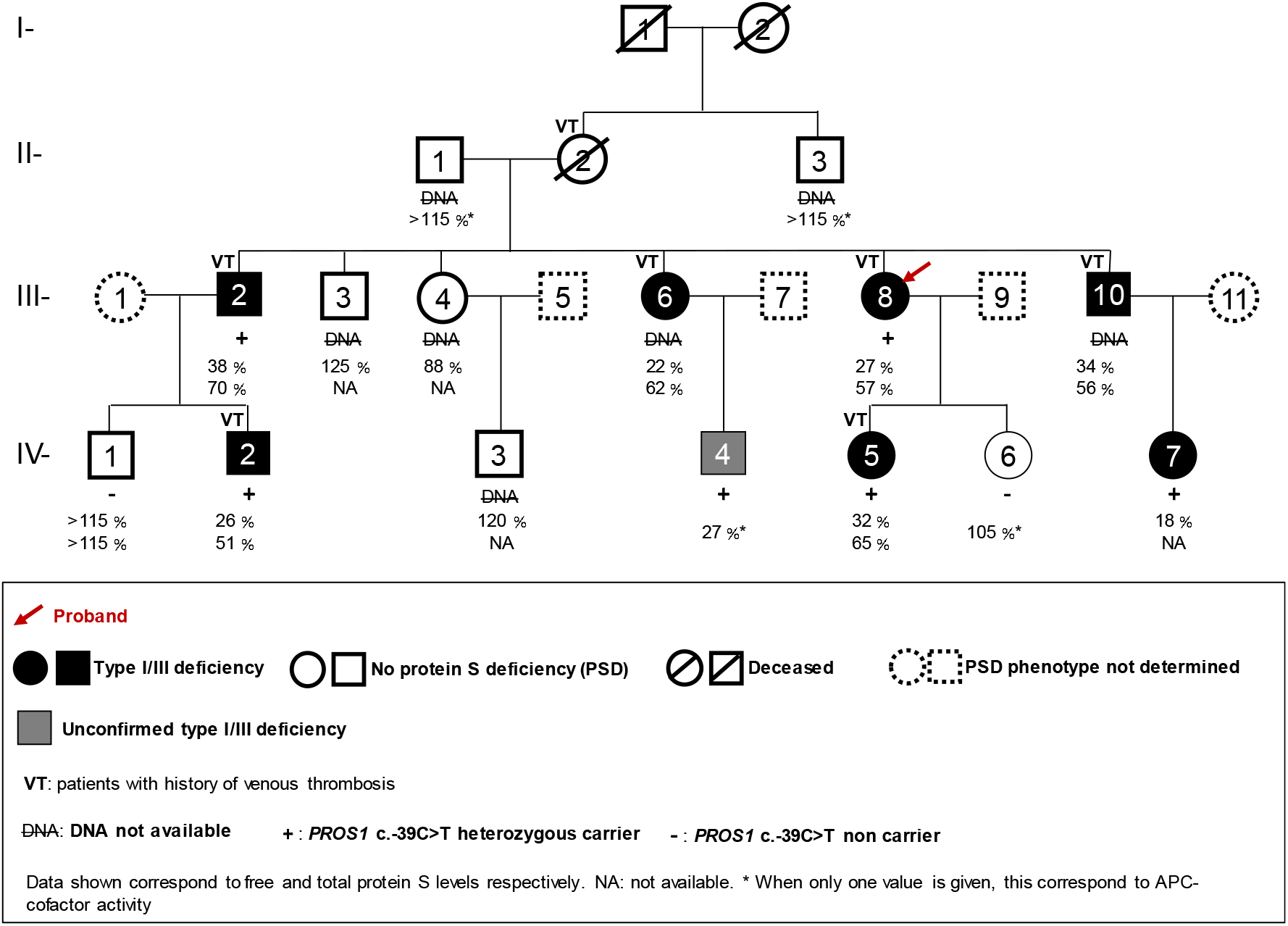
Pedigree of the studied family affected with Protein S deficiency.

Phenotypic protein S deficiency was diagnosed by measuring APC-cofactor activity, free (FPS) and/or total (TPS) proteins S levels. The cofactor activity of protein S was measured by an activated partial thromboplastin time-based clotting assay using the Staclot protein S kit (Diagnostic Stago, Asnieres, France) or by the IL test protein S kit (Instrument Laboratory Company, Milano,Italy). The concentration of total and free PS antigen were determined by an enzyme-linked immunosorbent assay (Asserachrom total or free protein S; Diagnostica Stago). In normal conditions, TPS levels range from 70% to 150% while normal FPS levels are generally greater than 60% in men, and 50% in pre-menopause women (>55% in menopause women). In the related PSD patients studied here, FPS levels ranged from 18% to 38% and TPS from 51% to 70% (Fig 1). APC-cofactor activity varied between 27% and 32%. Detailed clinical and biological information on family members are given in Supplementary Table 1.

All family members participating to this study signed a written informed consent for genetic investigations, according to the Helsinki declaration.

### Whole genome sequencing

With the aim of identifying the hypothesized culprit mutation causing the observed familial PSD, we sequenced the whole genome of 5 PSD patients (individuals III-2, III-8, IV-2, IV-4, IV-5) and 2 unaffected relatives (individuals IV-1 and IV-6) (Fig 1).

Whole genome sequencing was performed at the Centre National de Recherche en Génomique Humaine (CNRGH, Institut de Biologie François Jacob, Evry, FRANCE). After a complete quality control, 1μg of genomic DNA was used for each sample to prepare a library for whole genome sequencing, using the Illumina TruSeq DNA PCR-Free Library Preparation Kit, according to the manufacturer’s instructions. After normalisation and quality control, qualified libraries were sequenced on a HiSeqX5 instrument from Illumina (Illumina Inc., CA, USA) using a paired-end 150 bp reads strategy. One lane of HiSeqX5 flow cell was used per sample specific library in order to reach an average sequencing depth of 30x for each sequenced individual. Sequence quality parameters have been assessed throughout the sequencing run and standard bioinformatics analysis of sequencing data was based on the Illumina pipeline to generate FASTQ file for each sample. FastQ sequences were aligned on human genome hg37 using the BWA-mem program (Li & Durbin, 2009). Variant calling was performed using the GATK HaplotypeCaller (GenomeAnalysisTK-v3.3-0, https://software.broadinstitute.org/gatk/documentation/article.php?id=4148) tool followed by recalibration. Single nucleotide variants (SNVs) that succeeded the “PASS” filter were then annotated using Annovar (Wang *et al*, 2010).

To comply with the autosomal dominant mode of inheritance of the observed PSD, we selected as candidate culprit PSD causing mutation any rare variant that has either never been reported in public database or at a very low allele frequency (<1‰) and that were present at the heterozygous states in the 5 whole genome sequenced patients but not present in 2 healthy relatives. Interrogated public databases were dbSNP, GnomAD (https://gnomad.broadinstitute.org/) Ensembl (https://www.ensembl.org/index.html), 1000 genomes (https://www.internationalgenome.org/), ExAC (http://exac.broadinstitute.org/) and FrEx (http://lysine.univ-brest.fr/FrExAC/).

### Sanger sequencing validation

Sanger sequencing validation of the identified candidate mutation was carried out using primers designed to span all the putative transcription factor binding sites described by de Wolf (de Wolf *et al*, 2006) (Supplemental Table 2). PCR amplification of genomic DNA was performed using Phusion™ Green Hot Start II High Fidelity DNA Polymerase with GC Buffer in presence of 10% of DMSO (ThermoFisher). Big dye sequencing chemistry was used to sequence the PCR products in both directions using the ABI 3500xL Genetic analyser (Applied Biosystems, Foster City, CA) and the sequence were analyzed by the SeqScape Software (Applied Biosystems, Foster City, CA).

### Plasmid constructs

To assess the impact of the *PROS1* c.-39C>T variation, several expression vectors were investigated (Fig 2B). The human *PROS1* cDNA (GenBank NM_000313.4) was PCR-amplified from HepG2 cDNA and cloned into pcDNA3.1/*myc*-His(−) plasmid (Invitrogen). The PCR amplification of *PROS1* from c.-44 to c.2028 was performed using Phusion High Fidelity (HF) DNA Polymerase (ThermoFisher) and primer pairs designed to be either complementary to the wild-type sequence or carrying the c.-;39C>T point mutation on the forward primer only (Supplemental Table 3). After restriction enzyme digestion, the wild-type and mutant PCR products were cloned into pcDNA3.1 to fuse the full-length *PROS1* coding sequence in-frame with a myc-his tag, generating the wild-type (WT *PROS1*) and the mutant (Natural mutant *PROS1*) *PROS1* vectors, respectively (Fig 2B). To analyze the translation potential of the novel upstream start codon and overlapping uORF generated by the c.-39 C>T mutation, two additional expression vectors were constructed: (i) a truncated mutant *PROS1* vector was constructed by PCR-amplification of *PROS1* cDNA from c.-44 to c.110 (cloning of the overlapping uORF sequence) using mutagenic primers (Supplemental Table 3) fused in frame with the myc-His tag into pcDNA3.1; (ii) an elongated mutant *PROS1* vector (Elongated mutant *PROS1*) was obtained from the Natural mutant *PROS1* in which the new stop codon induced by the c.-39T allelic variant was suppressed by creating a 1-bp deletion (c.111delT) through rapid-site-directed-mutagenesis using phosphorylated non-overlapping primers (Supplemental Table 3) and Phusion HF DNA Polymerase (ThermoFisher), according to the manufacturer’s recommendations. All the recombinant plasmids were verified by Sanger sequencing (Genewiz).

**Figure 2:**
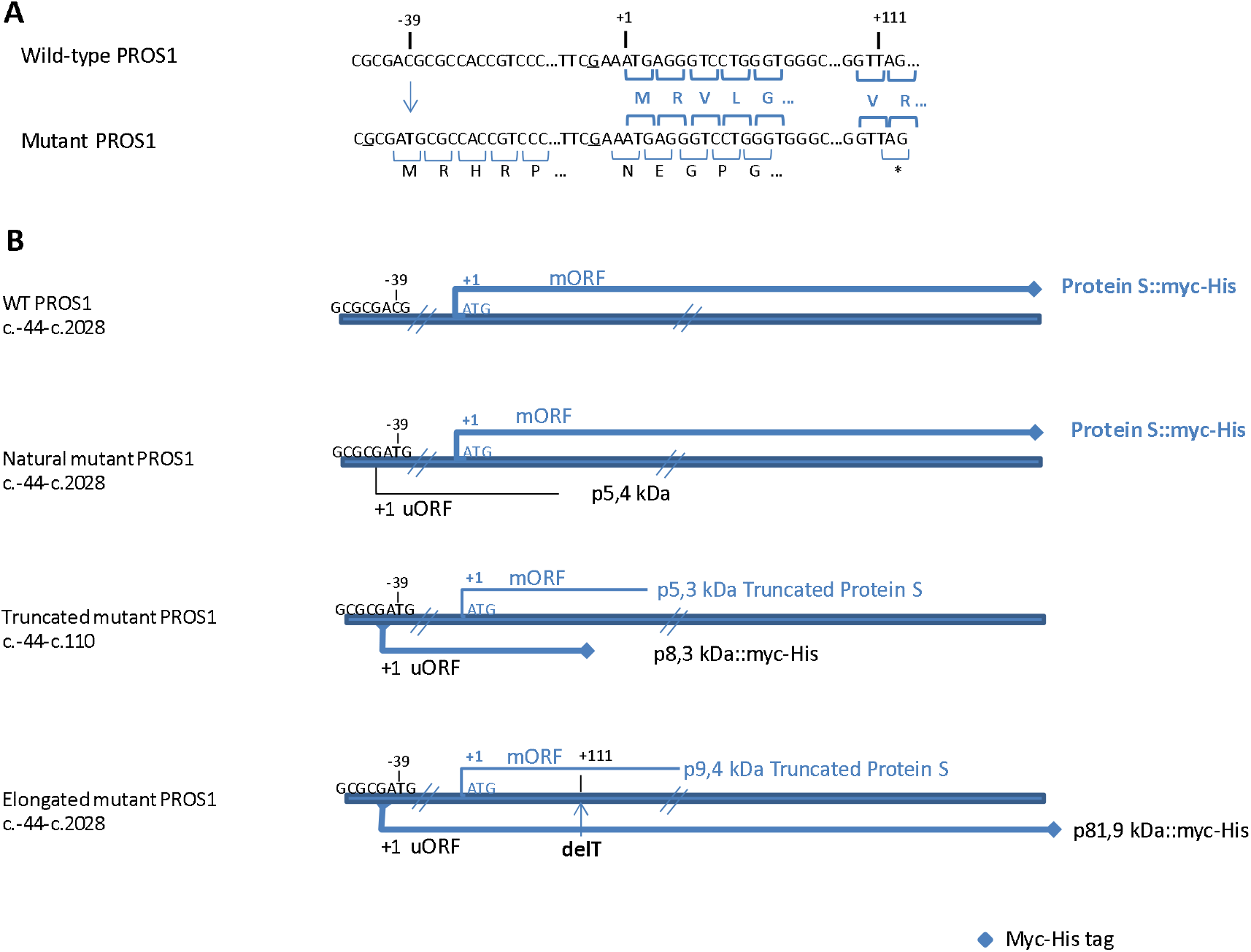
Schematic representation of the identified *PROS1* c.-39C>T mutation. (A) *PROS1* wild-type and c.-39C>T genomic sequences. Positions of the identified variant (c.-39), as well as the first nucleotide on the main ORF (+1) and of the generated stop codon (+111) are indicated above the WT sequence. Brackets indicate the main ORF and new uORF codons with the respective amino acid sequences. (B) The different molecular constructs used to investigate the mutation’s effect in vitro, as described under materials and methods.

### Cell culture and transfection

HeLa cells were cultured in RPMI medium (Gibco-Invitrogen) supplemented with 10% fetal calf serum (Gibco-Invitrogen), 1% penicillin/streptomycin and 1% Hepes. Cells were maintained in T75 flasks in a humidified atmosphere of 95% and 5% CO2 at 37°C. Twenty four hours before the transfection, 6-well plates of HeLa cells were prepared with 4×10 cells/well. Transfections were performed with jetPRIME® reagent (Polyplus Transfection) according to the manufacturer’s recommendations, with 0.5-1.5 μg/mL of each expression plasmid or empty pcDNA™3.1/*myc*-His(−) vector at a cell confluence between 60 and 80%. In some experiments, 0,5 μg of pcDNA3.1/*myc*-His/lacZ was included in each transfection to control the transfection efficiency. Cells were harvested and lysed 48 hours after transfection to extract total RNA and protein.

### RNA isolation and RT-qPCR analysis

Total RNA was isolated from the transfected HeLa cells using the RNeasy mini kit (Qiagen) following the manufacturer’s instructions. Reverse transcription of 2 μg of total RNA to single-stranded cDNA was performed using High Capacity cDNA kit (Applied Biosystems). To quantify *PROS1* transcripts, qPCR were carried out on the generated cDNA (diluted at 1/5) using PowerUp SYBR master mix (Thermofisher Scientific) in a final volume of 10 μl and 40 cycles of amplification on QuantStudio3 Real-Time PCR System (ThermoFisher). Results were analyzed with the QuantStudio design and analysis software. Transcript levels were normalized to the reference RPL32 gene. Relative quantification of *PROS1* in different samples was conducted according to threshold cycle (Ct) value based on the ΔΔCT method. All the primer pairs used in this study (supplemental Table 4) have a reaction efficiency between 90% and 110% and melting curves were analyzed to check the specificity of each RT-qPCR reaction.

### Protein preparation and western blot analysis

Protein extracts from the transfected HeLa cells were prepared by using 140 μL per well of RIPA buffer containing protease inhibitors. After clarification of the extracts by centrifugation, protein concentration was determined by the BCA method (Pierce™ BCA Protein Assay Kit). Equal amounts (50 μg) of total protein suspended in loading buffer were separated by SDS-PAGE using 8% or 4-20% gradient gels and transferred onto PVDF membranes (Immobilon-P or Immobilon PSQ, Merck Millipore) for immunostaining. Then, membrane was divided into two halves and probed with either monoclonal anti-(c-Myc) antibody (Merck Millipore) or anti-α-tubulin antibody (Sigma-Aldrich). After incubation with goat anti-mouse IgG Alexa Fluor 700 (ThermoFisher), blots were simultaneously scanned on Odyssey Infrared Imaging System (Li-Cor Biosciences) in the 700 channel. α-Tubulin was used as control.

### Statistical data analysis

Differential mRNA levels analysis according to experimental conditions were performed with ANOVA followed by Tukey’s HSD post-hoc tests for multiple comparisons. A statistical threshold of p <0.05 was used to declare statistical significance.

## RESULTS

With the aim of completely ruling out the possibility of PSD causing mutations in the structural gene, we first focused on genetic variations mapping the *PROS1* locus, downstream and upstream of the coding sequence. We thus identified a never reported c. -39C>T (chr3:93692632) mutation in the *PROS1* 5’UTR present at the heterozygous state in all 5 PSD sequenced patients and absent in the two sequenced healthy relatives. The rare T allele results into a novel upstream translation start codon (uATG) generating a novel overlapping upstream open reading frame (uORF) of 51 codons out-of-frame with the normal ORF (Fig 2A).

Because of the broad role of uORFs on downstream translation (Johnstone *et al*, 2016; Lin *et al*, 2019), we hypothesized that the detected mutation could be at the origin of the observed PSD. To validate this hypothesis, we first confirmed the presence of the mutation in the five sequenced patients by Sanger sequencing (Supplementary Fig 1) and genotyped it in one additional affected related (individual IV-7) for whom DNA was available. The absence of the the mutation in the two healthy relatives with available DNA was confirmed by Sanger sequencing as well.

Then, to experimentally assess the impact of the mutation on *PROS1* transcription and translation, we first generated 2 myc-His tagged molecular constructs (Fig 2B), the full length *PROS1* constructs containing either the wild-type ("WT") or the mutated -39T allele ("Natural mutant"). We observed that transfecting HeLa cells with the wild type or the natural mutant was homogeneously and significantly associated (p <10^-3^) with increased *PROS1* mRNA levels as compared to controls (Fig 3A and Supplemental Fig 2) without altering the expression of endogeneous *PROS1* (Fig 3B). Interestingly, the overexpression was not significantly different (p = 0.38) between WT and Natural mutant. These first results suggest that the c.-39C>T mutation does not impact the transcription machinery of the *PROS1* transgene.

**Figure 3:**
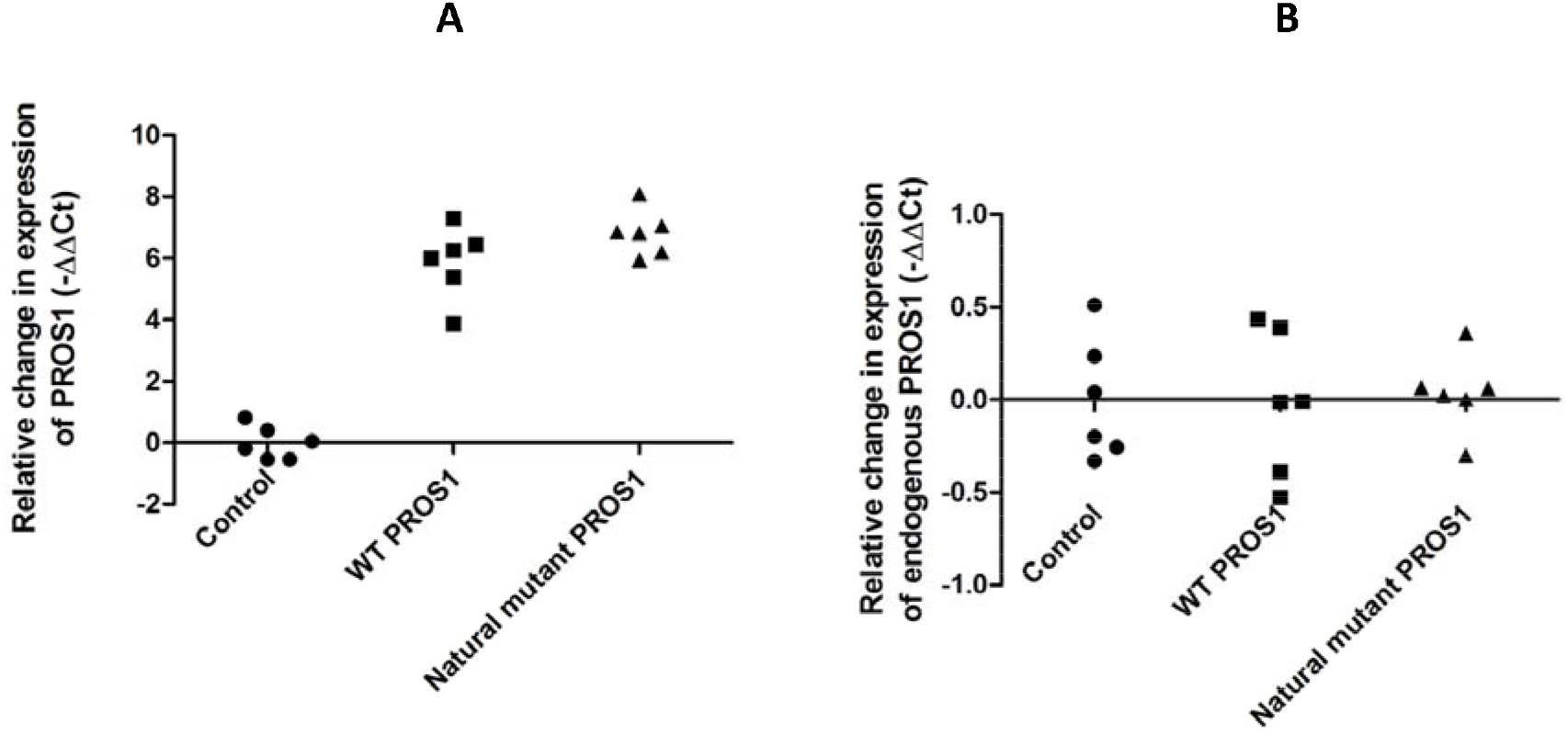
Impact of the *PROS1* c.-39C>T mutation on PROS1 RNA expression. mRNA expression levels were evaluated after transfection of HeLa cell lines with 1 μg of vectors encoding full-length WT PROS1 or Natural mutant PROS1. As control, cells were transfected with the empty pcDNA3.1/*myc*-His plasmid alone. After 48 hours of culture, mRNA expression was evaluated by Quantitative Reverse Transcription PCR (RT-qPCR) using primer pairs targeting either (A) total PROS1 expression (ex1fw,ex2rv primers) or (B) only the endogenous PROS1 (3UTRfw,3UTRrv primers). Results were normalized to RPL32 mRNA and analyzed by using the comparative Ct Method (ΔΔCt Method). -∆∆Ct Values are shown on y-axis.

We further investigated the synthesis of the fusion-myc-His construct proteins and confirmed that *PROS1* protein was produced in a dose-dependent manner by the wild type plasmid while no *PROS1* protein was detected with the natural mutant plasmid (Fig 4A). To better understand the underlying mechanisms, we generated the “truncated mutant” construct, containing the overlapping uORF in frame with the tag (Fig 2B). Our aim was to evaluate the ability of the novel upstream start codon to initiate the translation of an expected 8.3kDa fusion protein (https://www.snapgene.com/snapgene-viewer/). We failed to evidence any fusion protein synthesis in this molecular weight range (Fig 4B) while the expression of *PROS1* transgene was confirmed by qPCR (Supplemental Table 5). To go further, we generated an elongated mutant plasmid (see Materials and Methods) allowing the translation of a longer tagged protein from the identified upstream start codon. Western blot analyses showed the presence of a protein at the expected 81.9 kDa weight (Figure 4C), suggesting that a translation from the uAUG and driven by the uORF was initiated. However, the level of expression of the latter tended to be lower than that of the wild type protein, despite similar mRNA levels (Supplemental Table 5). This second set of experiments suggest that the c.-39C>T mutation creates a new functional uAUG codon that is able to initiate translation process while inhibiting the translation of the normal form of the Protein S from the main AUG.

**Figure 4:**
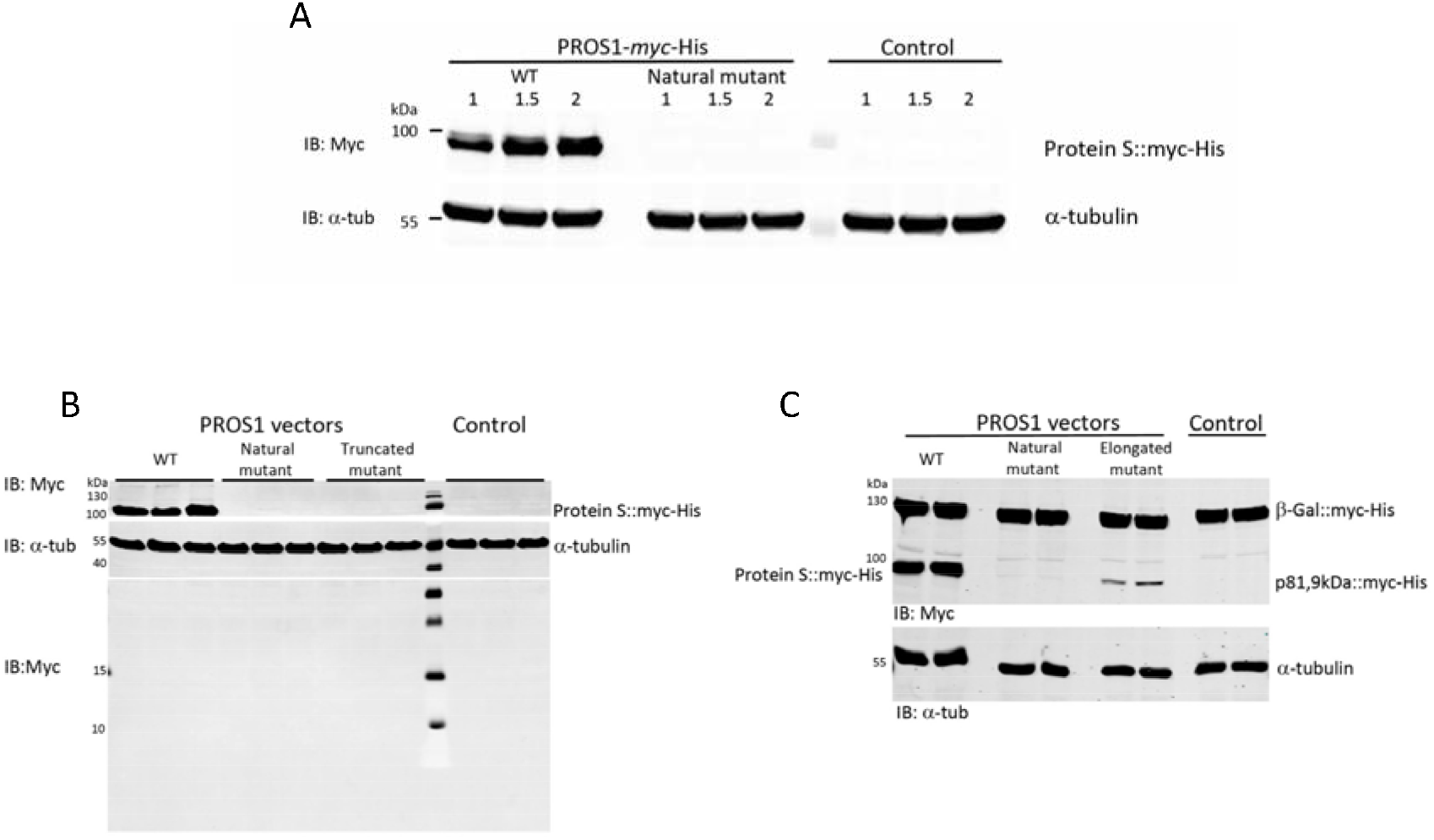
Impact of the *PROS1* c.-39C>T mutation on the Protein S expression. (A) Protein S expression was evaluated in HeLa cell lines transfected with 1, 1.5 and 2μg of either WT, Natural mutant PROS1 constructs or empty pcDNA3. 1/*myc*-His, as control (see Materials & Methods). Note that the wild-type construct directs expression of full-length Protein S in a dose-dependent manner, whereas this expression is blocked in the Natural mutant-PROS1 encoding the novel uAUG. B) Protein expression was evaluated in HeLa cell lines transfected with 1, 1.5 and 2μg of either WT, natural mutant PROS1, truncated mutant PROS1 or empty pcDNA3. 1/*myc*-His, used as control. The wild-type construct directs expression of the full-length Protein S. However, we have not detected any protein associated with the mutation (natural and truncated mutant PROS1 encoding the novel uAUG). (C) Protein S expression was evaluated in HeLa cell lines transfected with 1μg of vectors encoding WT PROS1, Natural mutant PROS1, Elongated mutant PROS1 or empty vector control. To control the transfection efficiency, 0,5 μg of pcDNA3.1/*myc*-His/lacZ was included in each transfection. Note that the wild-type construct directs expression of full-length protein S. Interestingly, we highlighted the presence of a larger mutant protein with the elongated mutant PROS1 construct, probably resulting from the use of the new AUG.

## DISCUSSION

We here report the first case of inherited PSD due to a rare mutation in the *PROS1* promoter region that creates a premature ATG codon out-of-frame relative to the natural ATG codon, thus generating a premature stop codon.

Following a negative target gene sequencing of the *PROS1* exonic sequences and large genomic rearrangement, a whole genome sequencing strategy was adopted to identify the molecular defect responsible for inherited PSD in a French pedigree with 8 PSD affected individuals and 2 unaffected relatives. This led to the identification of the novel c.-39C>T mutation, with the rare T allele generating a novel uAUG start codon and a novel overlapping uORF of 51 amino acids. Even if the flanking sequence of the new ATG does not completely match to the consensus predicted Kozak sequence ((G/A)NNAUGG), it seems similar to the one flanking the natural *PROS1* ATG, suggesting that the new ATG could be identified by the translational machinery and thus used in the cells (Kozak, 1997, 1990). Several works have previously demonstrated that uAUG and uORF are cis-regulatory elements in 5‘UTR that are able to control protein expression by altering the translation efficiency and could subsequently be associated with risk of diseases (Dvir *et al*., 2013; Lin *et al*., 2019; Orr *et al*., 2019; von Bohlen *et al*., 2017). Many previously ignored uORFs are now known to act as major post-transcriptional regulatory elements or to be translated to produce bioactive peptides or proteins (Jones *et al*, 2017; Orr *et al*, 2019).

Interestingly, the relative repression of uORF-containing mRNA was shown to be more pronounced when the uORF overlaps the translation starting site of the CDS (Johnstone *et al*, 2016). Our experimental data suggest that c.-39T allele does not impact on the *PROS1* transcription machinery but rather alters the translation of the main ORF leading to low Protein S production consistent with the observed PSD in carriers of the mutation. The c.-39T allele is expected to produce a shorter protein of 51 amino acids with no homology to wild-type Protein S. However, we were not able to detect it through our WB experiments. This prevented us from assessing whether the resulting protein is not stable, or not produced because of a possible degradation of the new mRNA by the nonsense mediated decay. Unfortunately, we were not able to collect mRNA materials from patients which also prevented us from investigating this hypothesis.

Importantly, none of the popular bioinformatics tools used to annotate variants such as PolyPhen, Sift, CADD (Rentzsch *et al*, 2019), TraP (Gelfman *et al*, 2017) or RegulomeDB (Boyle *et al*, 2012) were able to predict any deleteriousness for this mutation while the PreTIS software (Reuter *et al*, 2016) dedicated to 5’UTR variants predicted a “moderate” impact of the mutation.

While the identified c.-39C>T mutation segregates with PSD, it is also important to emphasize that all but one family members with PSD and complete clinical information experienced one or more VT events, with age of onset of first VT varying between 16 and 38 years old. A part from the PSD patient with missing clinical information (individual IV-4), the sole affected patient carrying the disease mutation that has not (yet) developed a thrombotic event is the youngest one. She is currently 16 yrs old and her free PS levels were 18%. Free PS levels in other relatives with combined PSD & VT ranged from 22% to 38% while such values ranged from 88% to 125% in healthy family members. As no thrombotic event was observed in non PSD relatives, these observations are in agreement with previous works suggesting genetically induced low free PS levels as a strong risk factor for VT. Indeed, Pintao *et al* observed that FPS levels lower than 33% was associated with an Odds Ratio (OR) for VT of ∼5 (Pintao *et al*, 2013). In a retrospective family, Lijfering *et al*. observed an increased OR for VT of ∼6 in relatives with FPS levels lower than 41% compared to relatives in the upper quartile of FPS distribution. Similarly, Alhenc-Gelas *et al* (Alhenc-Gelas *et al*, 2016) proposed to define a ∼30%-40% cut-off value for FPS levels as a way to identify individuals at high VT risk. All these proposed thresholds were consistent with the FPS levels observed in our studied family. As suggested by Alhenc-Gelas *et al*, the associated VT risk and the threshold value for FPS levels may be variable and may depend on the characteristics of the underlying causative variant. As an example, Suchon *et al*. demonstrated that the *PROS1* Heerlen mutation was associated with an OR for VT of ∼6 but a moderate decrease in free PS levels (∼ 70%)(Suchon *et al*, 2017).

Of note, this family project was set up in the context of the GenMed LABoratory of Excellence (http://www.genmed.fr/index.php/fr/), one of the objectives of which is to propose WGS in Mendelian disorders for which whole exome sequencing and target gene sequencing has been inconclusive. Even though WGS is now more and more affordable, it must be stressed that a less expensive deep sequencing of the *PROS1* locus including regulatory regions would have likely permit the identification of the culprit mutation. This illustrates the importance of exploring regulatory regions of known disease gene when searching for molecular diagnostics, a result that goes far beyond PSD but shall be generalized to other inherited (hematological) disorders.

## ACKNOWLEDGMENTS

O.S and M.R were financially supported by the GENMED Laboratory of Excellence on Medical Genomics [ANR-10-LABX-0013]. DA T was supported by the «EPIDEMIOM-VTE» Senior Chair from the Initiative of Excellence of the University of Bordeaux. This work was partially supported by the GENMED Laboratory of Excellence on Medical Genomics (ANR-10-LABX-0013) and the French Clinical Research Infrastructure Network on Venous Thrombo-Embolism (F-CRIN INNOVTE), two research programs managed by the National Research Agency (ANR) as part of the French Investment for the Future.

Clinical investigations of patients and of their relatives were conducted by SLM, CM, YH and JC. Whole sequencing was performed by CB, AB and RO. MR was in charge of the bioinformatics analysis of the whole genome sequenced data. Experimental investigations were conducted by CP, OS and BJV. The research study was designed by JFD, PEM, BJV and DAT. The manuscript was drafted by SLM, BJV and DAT and further reviewed by all co-authors.

