## Supplementary Data for "A novel rare c. -39C>T mutation in the *PROS1* 5’UTR causing PS deficiency by creating a new upstream translation initiation codon and inhibiting the production of the natural protein"

**SUPPLEMENTAL DATA**

**Supplemental Table 1**. Clinical characteristics of the pedigree members

| **Family characteristics** | | | | | | | | | | | | | |
| --- | --- | --- | --- | --- | --- | --- | --- | --- | --- | --- | --- | --- | --- |
| **ID** | **sex** | **Birth year** | **Age (yrs) at examination** | **VT episodes** | **Age at 1st VT (yrs)** | **PS Activity (**UI/dL**)** | **Free Protein S (**UI/dL **)** | **Total Protein S (**UI/dL **)** | **PS deficiency** | **Concomitant thrombophilic defects** | | | **NM_000313.3**  **PROS1 : c.-39C>T** |
|  |  |  |  |  |  |  |  |  |  | APC-R | FV:R534Q | FII:G20210A |  |
| II-1 | M | NA | NA | NA | NA | >115 | NA | NA | No | No | NA | NA | NA |
| II-2 | F | NA | Dead | 1 PE | 45 | NA | NA | NA | NA | NA | NA | NA | NA |
| II-3 | M | 1949 | 63 | 0 | - | >115 | NA | NA | No | No | NA | NA | NA |
| III-2 | M | 1950 | 59 | 1 DVT | 38 | NA | 38 | 70 | Type III | No | No | No | Heterozygote carrier |
| III-3 | M | 1953 | 62 | 0 | - | NA | 125 | NA | No | NA | NA | NA | NA |
| III-4 | F | 1956 | NA | 0 | - | NA | 88 | NA | No | No | NA | NA | NA |
| III-6 | F | 1960 | 56 | 1 SVT,  5 DVT^(1)^ | 16 | NA | 22 | 62 | Type I | No | NA | NA | NA |
| III-8 | F | 1962 | 46 | 4 DVT ^(2)^ | 20 | NA | 27 | 57 | Type I | No | No | No | Heterozygote carrier |
| III-10 | M | 1966 | NA | DVT+EP | 29 | NA | 34 | 56 | Type I | No | NA | NA | NA |
| IV-1 | M | 1974 | 35 | 0 | - | 146 | >115 | >115 | No | No | No | No | Non carrier |
| IV-2 | M | 1979 | 30 | 3 DVT, 2 EP ^(3)^ | 20 | NA | 26 | 51 | Type I | No | no | no | Heterozygote carrier |
| IV-3 | M | 1978 | NA | 0 | - | NA | 120 | NA | No | No | NA | NA | NA |
| IV-4 | M | 1987 | 13 | NA | NA | 27 | NA | NA | ambiguous | No | No | No | Heterozygote carrier |
| IV-5 | F | 1992 | 27 | 3 DVT ^(4)^ | 24 | 31 | 32 | 65 | Type I | No | No | No | Heterozygote carrier |
| IV-6 | F | 1998 | NA | 0 | - | 105 | NA | NA | No | No | No | No | Non carrier |
| IV-7 | F | 2002 | 16 | 0 | - | 32 | 18 | NA | Type I or III | No | No | No | Heterozygote carrier |

VT: Venous Thrombosis; DVT: Deep Vein Thrombosis; PE: Pulmonary Embolism; SVT: Superficial Vein Thrombosis; NA: Not Available (no plasma, no DNA or no clinical data available)

**^(1)^** 1 SVT at the age of 16 years followed by 5 DVT: 1 spontaneous episode at the age of 35 years, 1 post traumatic at 47 years and 3 post-partum.

^(2)^ 4 DVT: 3 spontaneous episodes at the age of 20, 38 and 39 years located at the leg and one at right arm after the placement of an intravenous line.

^(3)^ Recurrent episodes: 2 PE, the first at the age of 20 and the second at 26; surrounded by two idiopathic episodes at the age of 23 and 26 years

^(4)^ 2 SVT at the age of 24 years, followed by 1 DVP: spontaneous episode at the age of 26 years located in the veins of the feet

**Supplemental Table 2**. Primers used to amplify 5’-UTR region spanning all the putative transcription factor binding sites described by de Wolf

| **PromS-FW** | 5’-TCATTGACTTCCAGGTTTTGG-3’ |
| --- | --- |
| **PromS-RV** | 5’-CAGGACCCTCATTTCGAAGC-3’ |

**Supplemental Table 3**. Primers used to yield wild-type PROS1, natural mutant PROS1, elongated mutant PROS1 and truncated mutant PROS1 pcDNA3.1 constructs.

| PROSI wt_NheI fw | 5'-cgcgctagcgcgacgcgccacc-3' |
| --- | --- |
| PROSI_c.-39C>**T**_ Nhe1 fw | 5'-cgcgctagcgcga**t**gcgccacc-3' |
| PROSI wt_Not1 rv | 5'-cgcgcggccgcagaattctttgtctttttccaaactg-3' |
| Mutant PROS1 fs delT fw | 5’P-AGGAAGCGTCGTGCAAATTCTTTAC-3’ |
| Mutant PROS1 fs delT rv | 5’P-ACCAGGACTTGTGAAGCCTGTTG-3’ |
| Mutant partial PROS1_Not1_ c.-39C>**T** fw | 5'-tagcggccgcgcga**t**gcgccacc-3' |
| Mutant partial PROS1_HindIII rv | 5'-gtaaagcttaccaggacttgtgaagcctg-3' |

**Supplemental Table 4**. Primers used in qPCR analyses.

| qPCR-PROS1 ex1fw | 5'-GAAATGAGGGTCCTGGGTGG-3' |
| --- | --- |
| qPCR-PROS1 ex2rv | 5'-CCAGGACTTGTGAAGCCTGT-3' |
| qPCR-PROS1 ex1fw | 5’-CTTCGAAATGAGGGTCCTGGG-3’ |
| qPCR-PROS1 ex1rv | 5’-GAGACGGGAAGCACTAGGAG-3’ |
| RPL32 Fw | 5'-CCCAAGATCGTCAAAAAGA-3' |
| RPL32 Rv | 5'-TCAATGCCTCTGGGTTT-3' |
| qPCR-PROS1 3UTRfw | 5’-GTGTGGGTCACACAAGGTCT-3’ |
| qPCR-PROS1 3UTRrv | 5’-CAGGCACGTGGCAATCTTAC-3’ |

**Supplemental Table 5**. Relative expression of PROS1 transgene in HeLa cells transfected with PROS1 mutants

| −ΔΔCt^a^ | | −ΔΔCt^b^ | |
| --- | --- | --- | --- |
| Control | Truncated mutant | Control | Truncated mutant |
| 0.82 | 11.84 | 0.78 | 12.46 |
| 0.05 | 11.72 | 0.27 | 11.55 |
| 0.40 | 11.26 | 0.45 | 11.35 |
| -0.54 | 8.27 | -1.71 | 9.06 |
| -0.55 | 10.95 | 0.02 | 11.44 |
| -0.18 | 11.48 | 0.20 | 12.61 |

| −ΔΔCt^a^ | | −ΔΔCt^b^ | |
| --- | --- | --- | --- |
| Control | Elongated mutant | Control | Elongated mutant |
| 1.29 | 11.64 | 0.15 | 10.19 |
| 1.35 | 11.26 | 0.57 | 10.68 |
| 0.85 | 10.93 | 0.52 | 10.26 |
| -0.98 | 10.34 | -0.31 | 9.40 |
| -1.15 | 10.89 | -0.48 | 9.55 |
| -1.36 | 11.07 | -0.46 | 11.27 |

^a^ qPCR results obtained using the (ex1fw,ex1rv) pair of primers

^b^ qPCR results obtained using the (ex1fw,ex2rv) pair of primers

**Supplemental Fig 1**. Screenshot of the Sanger sequencing validation of the presence of the c.-39C>T variant at the heterozygous state in carriers. The obtained peaks at the variant position are framed.

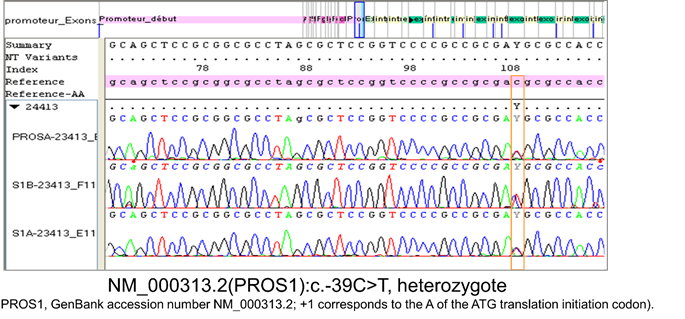

**Supplemental Fig 2**. Impact of the *PROS1* c.-39C>T mutation on PROS1 mRNA levels quantified by the (ex1fw, ew1rv) pair of primers. The relative change in PROS1 expression (−ΔΔCt) is shown.

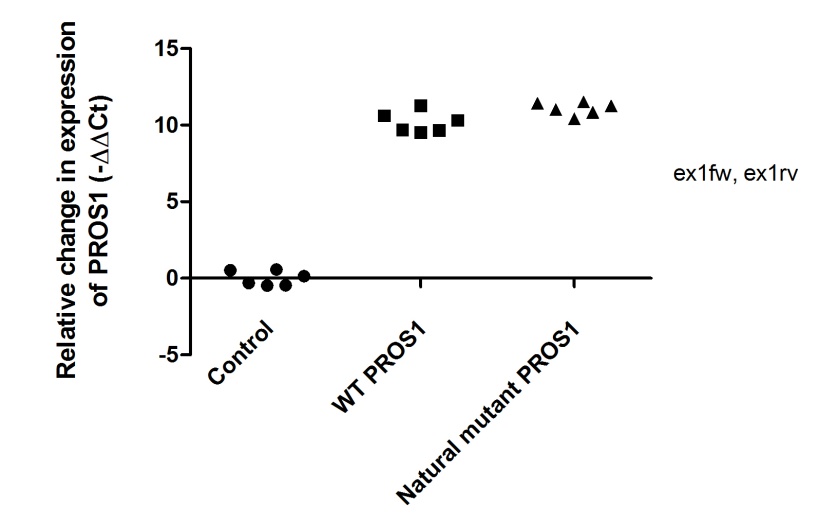
